# Modeling changes in vascular and extracellular matrix integrity that can occur in decellularized kidneys after implantation

**DOI:** 10.1101/2021.02.09.430397

**Authors:** Peter R. Corridon

**Author notes:** **Contact Information**: Peter R. Corridon, PhD, Assistant Professor, Department of Immunology and Physiology, College of Medicine and Health Sciences, Director, Pre-Medicine Bridge Program, College of Arts and Sciences, Khalifa University of Science and Technology, PO Box 127788, Abu Dhabi, UAE, Office.

## Abstract

A method was established to identify alterations in vascular patency and extracellular matrix integrity of decellularized porcine kidney scaffolds. These scaffolds were perfused with blood at physiologically normal (500 and 650 ml/min) and abnormal (200 ml/min) rates. Variations in venous outflow were then assessed over 24 hours. Angiographic data confirmed that standard arterial branching patterns and the integrity of the extracellular matrix were considerably disrupted. Scaffolds subjected to normal arterial perfusion rates observed drops in venous outflow across the 24 hours. These reductions rose from roughly 40% after 12 hours to 60% after 24 hours. At the end of the test period, regardless of the underlying damage that occurred, the kidneys appeared intact on the surface, and there were no apparent signs of clotting. In comparison, venous flow rates decreased by 80 to 100% across the 24 hours in acellular scaffolds subjected to a far lower perfusion rate of 200 ml/min. These kidneys also appeared intact after 24 hours of perfusion, but presented several arterial, venous, and ureteral clots. The results of this study provide insight into circumstances that limit scaffold viability and provide a simplified model to analyze other conditions that can better prepare scaffolds for long-term transplantation.

## Introduction

Nearly a billion individuals worldwide are affected by acute and chronic kidney conditions [1]. Presently, there are no cures for either acute kidney injury (AKI) or chronic kidney disease (CKD), and the prevalence of these debilitating diseases is on the rise. The escalating incidences of both conditions produce overwhelming burdens on healthcare systems, and studies have reported high rates of transition from AKI to CKD [2]. These renal complications support progressive and irreversible damage that often leads to end-stage renal disease (ESRD) [3]. This devastating disease progression illustrates a growing public health issue that results in high morbidity and mortality rates [4], and highlights the need to improve kidney disease management.

Once a renal disease progresses to ESRD, the kidneys can no longer perform their normal functions. At that point, clinical options are limited to renal replacement therapy (RRT), which consists of various treatment modalities that replace the normal filtration, secretion, reabsorption, endocrine, and metabolic functions of the kidney in varied capacities. These modalities are often grouped into two main categories: dialysis and transplantation [5]. While dialysis techniques replace lost filtration capacities, which help remove toxins and regulate blood pressure and pH, transplantation is the ultimate solution to reinstate all innate kidney functions. Unfortunately, the severe global shortage of transplantable kidneys [6], as well as organ rejection [7] limit this ideal option and accentuate the demand for alternative solutions.

Fortunately, recent advances in tissue engineering and regenerative medicine may help address this need. Such advances support the development of bioartificial kidneys, which can potentially increase the number of available transplantable kidneys, reduce transplant rejection rates, and significantly diminish morbidity and mortality in patients with acute and chronic disorders [8]. One viable pathway to developing transplantable devices relies on whole-organ decellularization to create natural scaffolds [9]. Researchers use this technique to create scaffolds by isolating the intact extracellular matrix (ECM) from human [10], canine [11], ovine [12], and porcine kidneys [13]. Acellular scaffolds then provide a platform for cell growth, differentiation, as well as tissue and organ development.

In the last few years, significant efforts have been made to optimize whole kidney decellularization techniques. Studies conducted on porcine kidneys have provided promising results with perfusion-based protocols using detergents [14]. For instance, the combination of Triton X-100 and sodium dodecyl sulfate (SDS) can efficiently remove cellular components and preserve ECM as well as vascular architectures, and has been identified as a reliable method to decellularize whole kidneys [14]. A study using this decellularization procedure has examined the vascular patency of acellular kidney scaffolds within the first two hours after transplantation [15]. The results of this study indicated that scaffolds maintained vascular patency only in the short-term under extremely low and non-physiologically relevant blood perfusion rates. Whereas, other studies have shown that decellularized porcine kidney scaffolds implanted in pigs developed thrombosis of the entire vascular tree, while withstanding physiological blood pressures for two weeks in vivo [16]. These results have provided crucial insight, but there is still a limited understanding of decellularized scaffold integrity, in general, and specifically, in post-transplantation settings [17].

The current study aimed to investigate whether alterations in decellularized scaffolds, created from the above combination of detergents, can be modeled in vitro to assess post-transplantation settings. To address this aim, a bioreactor system was created to subject scaffolds to continuous perfusion of whole blood at various rates over 24-hours. It was hypothesized that this model could be used to identify changes in the patency of the vasculature and integrity of the ECM. The results of this study would provide insight into circumstances that limit scaffold viability and provide a simplified model to analyze other conditions that can better prepare scaffolds for long-term transplantation.

## Results

### Assessment of Vascular Patency and ECM Integrity

The entire decellularization process produced acellular kidney scaffolds within 144 hours (6 days). The progressive changes in the renal structure that occurred are outlined in Figure 1. The data collected in Figure 2 illustrate intact renal vasculature and ECM of the native porcine kidney (Figure 2c), which were not disrupted by decellularization.

**Figure 1.**
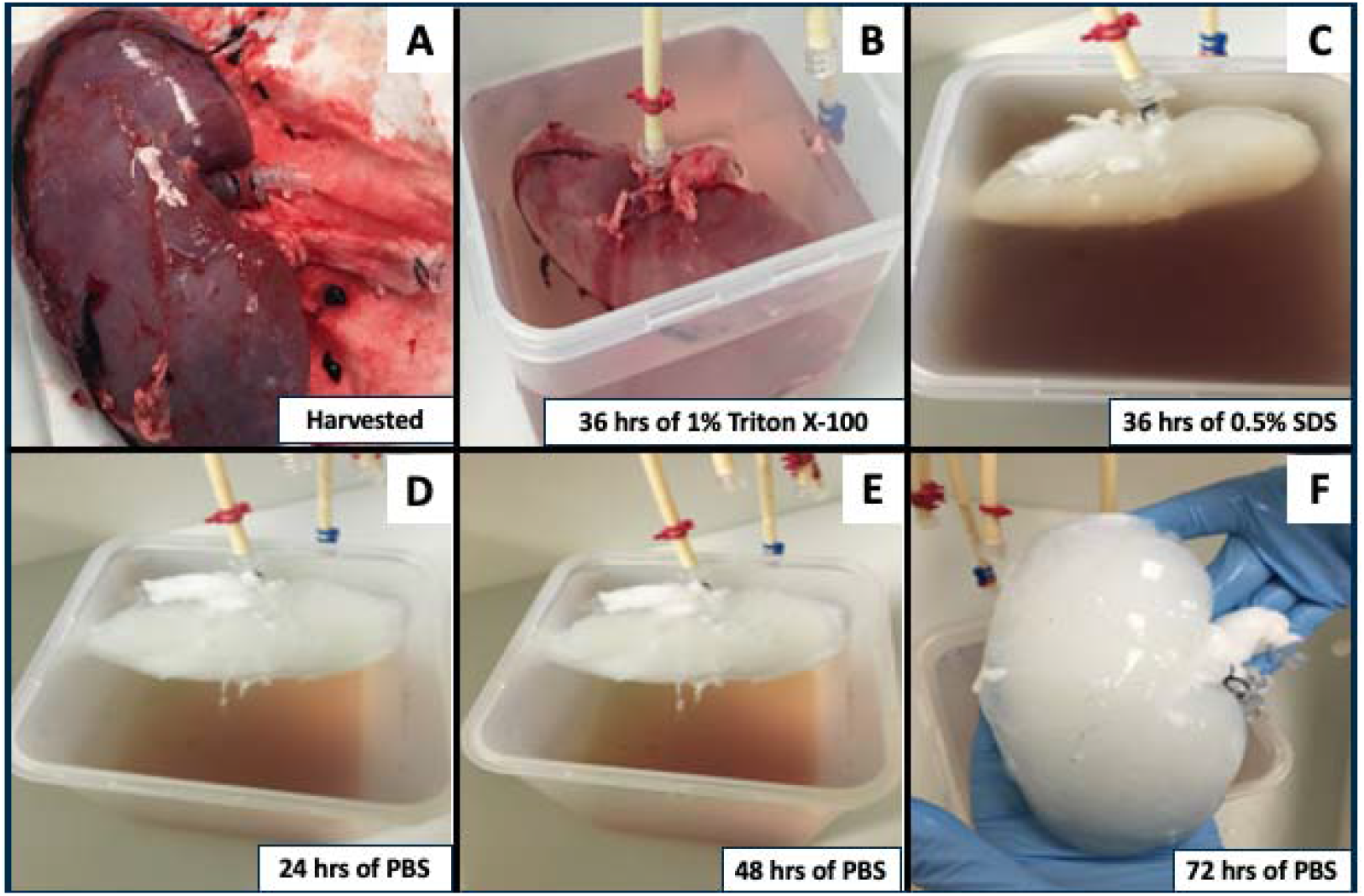
Generating whole kidney scaffolds using perfusion-based decellularization. (A) Kidneys that were harvested from adult Yorkshire pigs were cannulated and then subjected to the slow infusion of a sequential combination of detergents. These images show the results after treatment with Triton X-100 (B) and SDS (C), and PBS (D through F), and how this perfusion-based protocol successfully removed intrinsic cellular components to produce the finished decellularized product.

**Figure 2.**
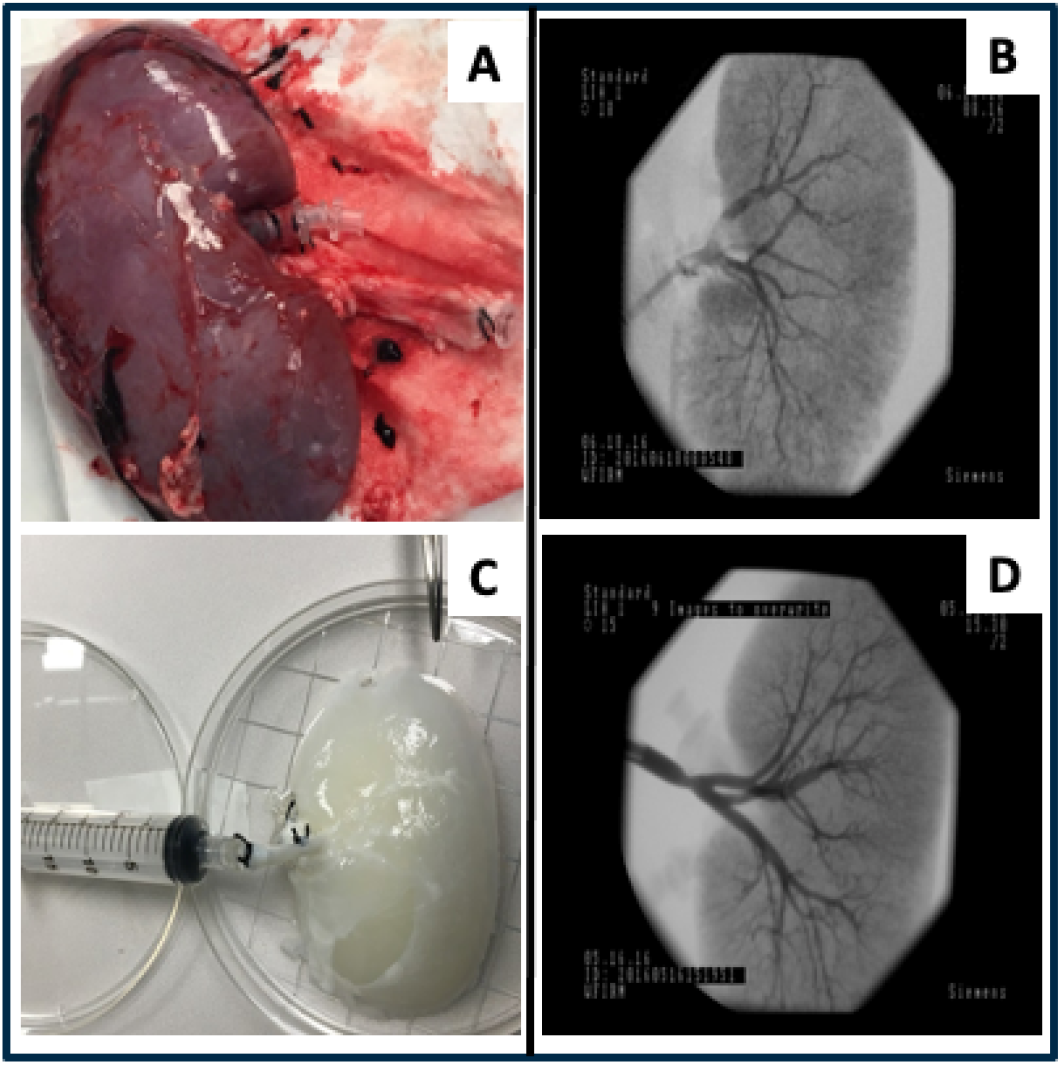
Preserved vascular and ECM architectures after perfusion-based decellularization. Whole native (A) and decellularized (C) porcine kidneys were infused with the contrast agent, and angiograms were collected to display intact vascular and ECM architectures of the respective organs (B and D). Note the pictures are of two different kidneys and are used for illustrative purposes only.

### Structural Changes in Scaffold Vascular and ECM Architectures

Angiographic images of decellularized kidneys showed clear visualization of the main renal artery, as well as, segmental, lobar, and interlobar arteries (Figures 3A, 3B, and 3C). Once again, these images illustrate how the vasculature architectures in these scaffolds were maintained after decellularization, and at the beginning of the blood perfusion studies. However, standard arterial branching patterns were considerably disrupted by the end of the study (Figures 3D, 3E, and 3F).

**Figure 3.**
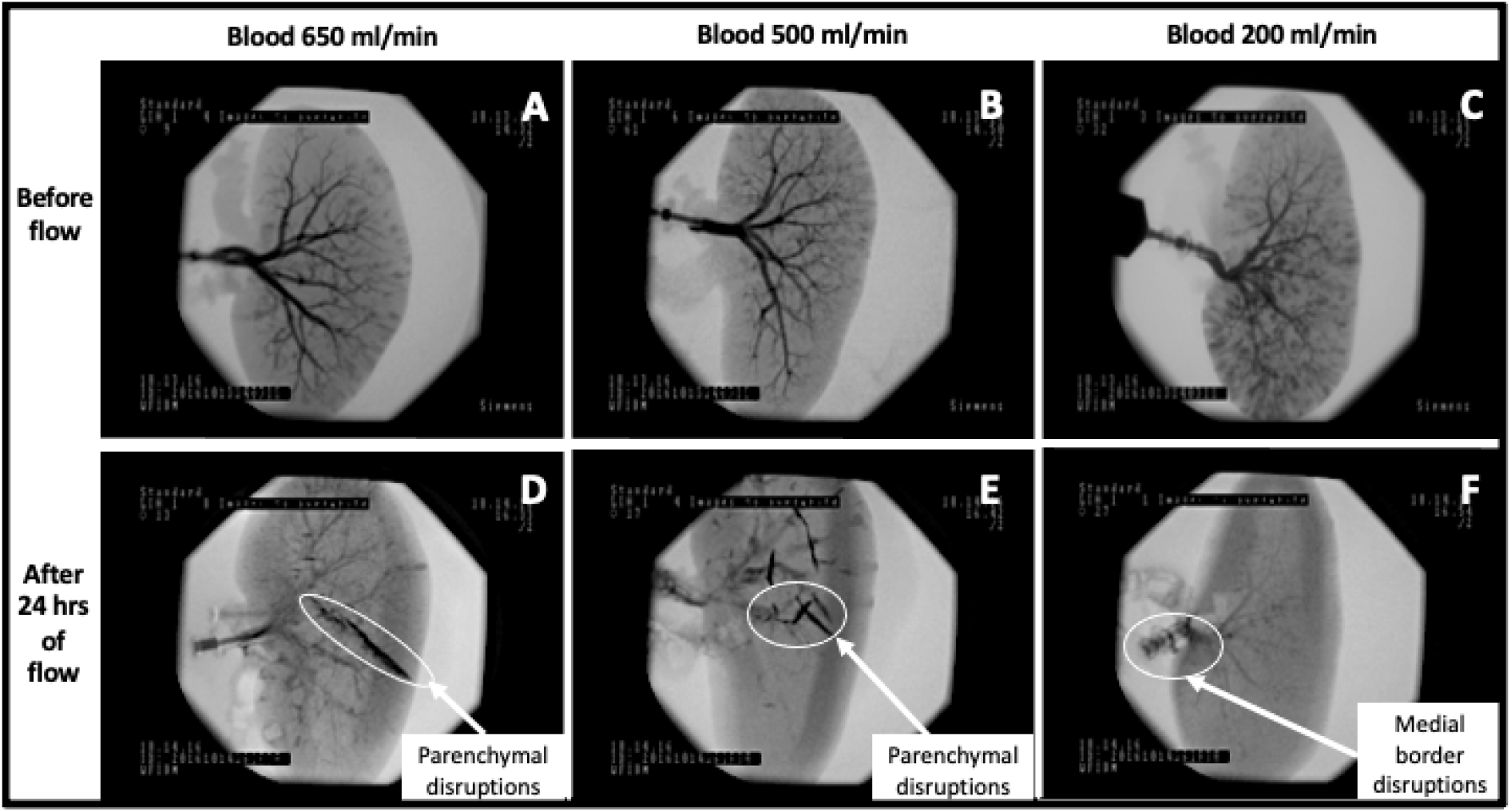
Comparisons between angiograms taken before and at the end of the blood perfusion studies. Images (A) through (C) outline intact vascular and ECM structures prior to whole blood perfusion. In comparison, angiography data reveal significant changes in the vascular architecture that occurred with blood perfusion (D, E, and F). The higher flow rates also appeared to generate visible parenchymal damage (E and F).

After 24 hours of hypothermic blood perfusion, the acellular kidneys that were perfused at normal arterial inflow rates were unable to maintain a standard vascular architecture. Furthermore, these changes in vascular patency were accompanied by substantial damage throughout the parenchyma that was not visible from the surface of the scaffold. These vascular networks were impaired in different ways. Specifically, there were instances when the scaffold integrity was compromised more in the lower pole than in the upper pole and vice versa (Figure 3E). Whereas, the kidneys that received 200 ml/min arterial inflow showed far lower signs of an ability to perfuse blood throughout their vascular networks. In such kidneys, there were more notable signs of vascular damage around the medial borders (Figure 4F) that resulted in subsequent impairment throughout the vasculature.

**Figure 4.**
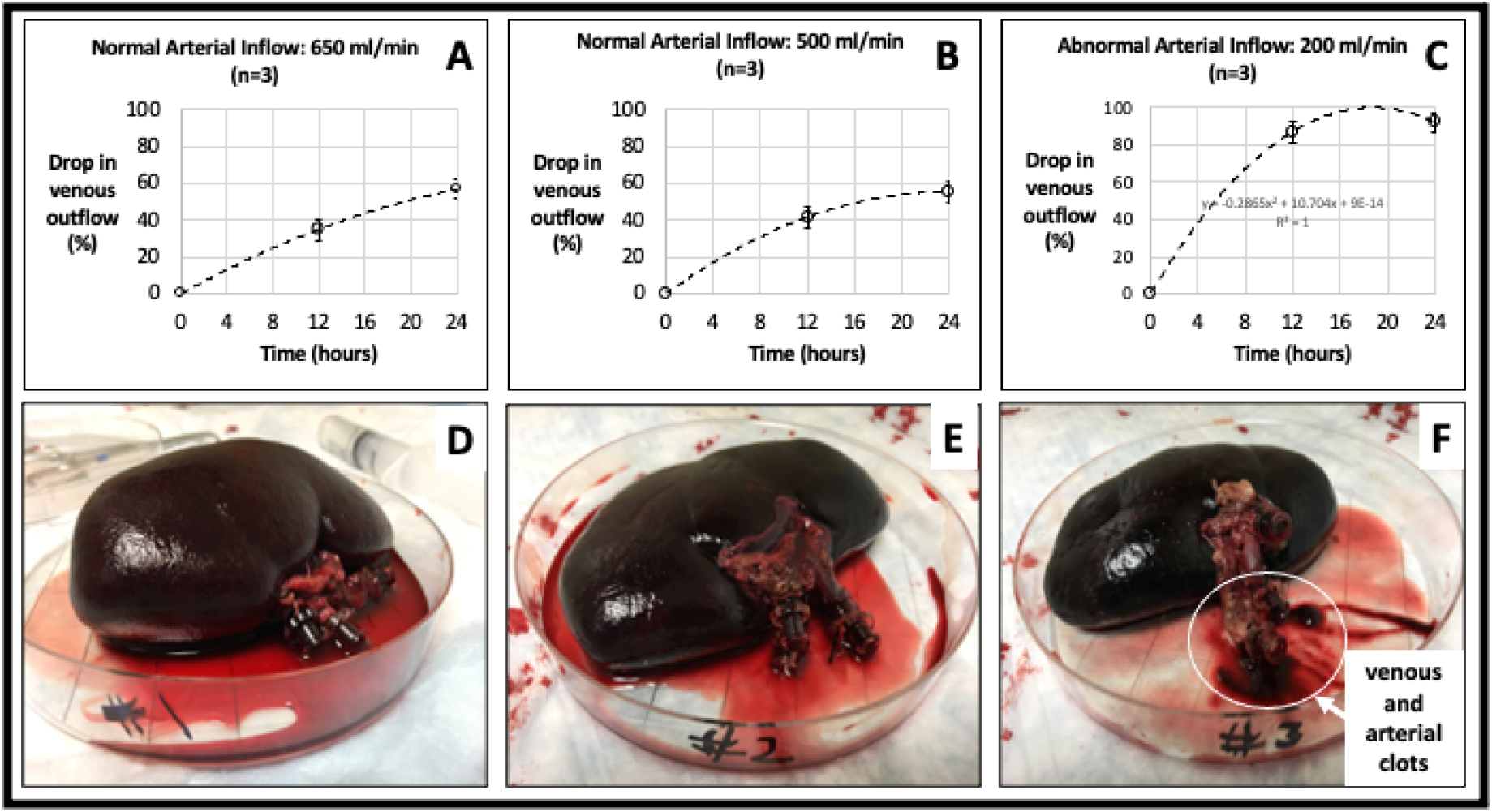
Reductions in venous blood outflow. The graphs in (A), (B), and (C) show the substantial, yet varied reductions in venous outflow as a function of arterial inflow and time. Image (F) highlights the conditions that supported the development of visible arterial, venous, and ureteral thrombi within the 24-hour timeframe, as compared to the conditions associated with images (D) and (E).

### Alterations in Scaffold Venous Outflow

At the beginning of each perfusion study, a steady flow of blood emerged from both the renal vein and ureter. However, as perfusion continued, there were significant reductions in blood outflow throughout the test period, as shown in images 4A, 4B, and 4C. The decellularized kidneys that received typical arterial perfusion rates (500 and 650 ml/min) had far lower reductions in venous output when compared to the scaffolds that received an abnormally low arterial perfusion rate (200 ml/min).

With arterial perfusion rates of 500 and 650 ml/min, there were gradual drops in venous outflow across the 24 hours. These reductions rose from roughly 40% after 12 hours to 60% after 24 hours (Figures 4A and 4B). At the 24-hour mark, these kidneys still perfused blood that was continuously recirculated and unreplenished. Specifically, the decellularized organs were still able to successfully expel nearly 40% of unreplenished blood after 24 hours. And, at the end of the test period, the kidneys appeared intact, and there were no apparent signs of clotting (Figures 4D and 4E).

In comparison, the acellular scaffolds that were subjected to a far lower perfusion rate produced much sharper reductions in the venous outflow. These scaffolds were poorly perfused as their venous flow rates decreased by 80 to 100% across the 24 hours (Figure 4C). Similarly, these hypoperfused kidneys also appeared intact at the end of the study but presented several arterial, venous, and ureteral clots (Figure 4F).

The Student’s t-test revealed no significant difference between the mean reductions in venous outflow that were observed with arterial perfusion rates of 500 and 650 ml/min (p = 0.439). However, the drops in venous outflow that occurred with arterial perfusion rates of 500 ml/min and those associated with a 200 ml/min flow rate were significantly different (p = 0.0147). There was also a significant difference between mean reductions in renal outflow obtained from 650 ml/min and 200 ml/min flow rates (p = 0.0298). For all arterial infusion rates, the venous outflow rates were also significantly different at the beginning and end of the perfusion studies (flow rate = 650 ml/min, p = 0.004; flow rate = 500 ml/min, p = 0.003, and flow rate = 650 ml/min, p = 0.002).

## Discussion

The current study aimed to investigate whether alterations in decellularized scaffolds can be modeled in vitro to assess post-transplantation settings. The decellularization process produced outcomes consistent with previous studies [18], and thus provided a means to investigate the macroscopic architectures of whole organ scaffolds. The results showed that substantial disruptions to vascular patency and ECM integrity occurred with renal blood flow variations across the 24-hours. Such continuous blood flow significantly reduced the structural and functional capacities of the acellular organs. More interestingly, kidney scaffolds perfused with unreplenished blood at physiologically normal rates were still capable of perfusing blood at the end of the study.

The steady flow of blood that emerged from both the renal vein, at the beginning of each experiment, signified that each arterial infusion flow rate supported whole organ perfusion. Thus, it was possible to initially utilize the intact arterial tree within the scaffold to deliver blood throughout the organ. Nevertheless, as perfusion continued in all cases, there were significant reductions to renal vein blood flow, even though arterial renal blood flow was kept constant.

Renal blood flow is roughly 20 to 25% of the cardiac output, ranging from 4 to 8 L/min [19]. The arterial inflow rates of 500 and 650 ml/min used in this study correspond to the amount of blood each kidney would receive during resting conditions. Under normal conditions, the kidneys autoregulate renal blood flow to ensure that pressure elevations are not transmitted to glomeruli or capillaries. Decellularized whole kidneys are incapable of autoregulation, and thus our experimental design compensated for this loss of function. The consistent supply of typical systolic renal blood helped examine how the scaffolds respond in post-transplantation settings.

By lowering the inflow rate to 200 ml/min, renal blood pressure was substantially reduced to approximately 25 mmHg. This hypoperfused state helped mimic arterial stenosis, which is a leading cause of graft failure [20]. The model did not fully mimic all aspects of transplant renal artery stenosis observed in vivo, since the scaffolds could not inherently generate compensatory increases in pressure. However, recirculating unreplenished blood provided an additional means to produce vascular occlusions that could have increased blood pressure, and ultimately facilitated damage to glomerular and capillary structures throughout the scaffolds. It is also feasible that the higher and continuous perfusion rates would have supported the development of intrarenal pressures that were detrimental to the microvasculature. However, the higher flow rates may have been sufficient to clear enough cells and tissue debris to maintain higher venous outflow levels by the 24-hour mark.

According to Hagen-Poiseuille’s law, renal perfusion pressure is a very important parameter to support blood flow and could also be used to explain the observed modifications in vascular structure. This law states, 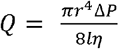 describes how blood flow (*Q*) can be modeled as a function of blood pressure (Δ*P*), viscosity (*η*), vessel length (*l*), and radius (*r*). From this law, hypothermic perfusion and reductions in arterial blood would have facilitated increases in blood viscosity [21]. Moreover, as the blood was not replenished, the various blood cells, in particular, the red blood cells, were continuously exposed to low glucose and oxygen environments. These conditions would have supported hemolysis [22], eryptosis [23], and aggregation [24], and further increases in blood viscosity with time. Such compounded increases in blood viscosity could have, in turn, supported renal artery stenosis and ischemia [25].

Compared to erythrocytes, leukocyte, and platelet activities, which rely on mitochondrial machinery to meet normal energetic demands, would have also been impacted by this deficiency in nutrients. Mitochondria dysfunction that would have resulted from hypoxia is known to diminish platelet survival severely and again increase the likelihood of vascular thrombosis [26], and leukocyte sequestration [27]. It has also been established that the decellularization process can reduce the mechanical strength of the ECM [28], and diminish the critical support needed to maintain the vasculature. Moreover, scaffold remnants have also been shown to recruit blood cells and influence an immune response in decellularized organs and [29], which can contribute to ECM damage.

The continuous recirculation of unreplenished blood would have also created acidic environments within the scaffolds, which could have again supported further degradation of sensitive ECM structures [29]. This process could have furthered attenuated the mechanical integrity of the scaffolds. Thus, the combinations of these events would have led to the overall impairment of both the ECM and vascular architectures generated from this model.

## Conclusion

The original and direct empirical evidence provided by this study illustrates how decellularized scaffolds may perform after implantation. This bioreactor model provides a means to investigate the time-based alterations in vascular and ECM integrities. It was shown that arterial blood flow and pressure have important effects on scaffold viability. In the future, it would be of interest to consider whether replenishing blood and heparin levels throughout the study could increase the functional capacity of the scaffolds and improve long-term transplantation.

## Materials and Methods

### Porcine Kidney Perfusion-based Deellularization and Sterilization

Adult Yorkshire pigs were euthanized under the guidelines provided by the Institutional Animal Care and Use Committee (IACUC) at the School of Medicine, Wake Forest University. All experimental protocols complied with all relevant ethical guidelines and regulations provided by this IACUC and the Animal Research Oversight Committee of Khalifa University of Science and Technology to ensure that animals were treated ethically and humanely. Moreover, all methods were performed in accordance with the ARRIVE guidelines. Whole porcine kidneys, with intact renal arteries, veins, and ureters, were decellularized and sterilized using methods outlined in detail in the literature [18]. Decellularization occurred in a non-sterilized environment. A sequential combination of detergents (Triton X-100 and SDS) and phosphate-buffered saline (PBS) was slowly infused into cannulated renal arteries, each at a rate of 5 ml/min. First, 1% Triton X-100, which was dissolved in deionized water, was perfused through the renal artery for 36 hours. Second, 0.5% SDS in PBS was infused for an additional 36 hours. Last, the kidneys were perfused with PBS for 72 hours to remove residual traces of detergents and cellular components. After decellularization, the scaffolds were submerged in PBS and then sterilized with 10.0 kGy gamma irradiation[13].

### Fluoroscopic Angiography

Whole native and decellularized kidneys were first flushed with PBS (approximately 100 ml) via the renal artery. Iothalamate meglumine contrast agent (60% Angio-Conray, Mallinckrodt Inc., St Louis, MO, USA) was then infused into the artery. Once a sturdy flow of contrast exited the renal vein, the renal vein, renal artery, and ureter were occluded to prevent contrast agent from leaking out the organ. Angiograms were conducted using a Siemens C-Arm Fluoroscope (Siemens AG, Munich, Germany) to investigate macrovascular patency and ECM integrity.

### Blood Perfusion Studies

Bioreactor components were sterilized using a ^60^Co gamma Irradiator: suction pump heads (Ismatec, Cole-Palmer, Wertheim, Germany); standard pump tubing female and male luer x 1/8፧ hose barb adapters; barbed fittings; reducing connectors; and kynar adapters (Cole-Palmer, Vernon Hills, IL, USA). The bioreactor tubing, chambers, and 2000 ml round wide mouth media storage bottles with screw cap assemblies (Sigma-Aldrich, St. Louis, MO, USA) were all sterilized using an autoclave.

After sterilization, bioreactors were assembled within a biosafety cabinet, and prepared for perfusion studies, as shown in Figure 5. Further details on the bioreactor are presented elsewhere [13]. Scaffolds were separated into three groups according to the blood perfusion rate: kidneys in group 1 (n=3) were perfused at a rate of 650 ml/min; kidneys in group 2 (n=3) were perfused at a rate of 500 ml/min; and kidneys in group 3 (n=3) were perfused at a rate of 200 ml/min. Prior to hypothermic perfusion, scaffolds were suspended in approximately 400-500 ml of heparinized pig whole blood (BioIVT, Westbury, NY, USA). The Ismatec MCP-Z Process or MCP-Z Standard dispensing pumps (Cole-Palmer, Vernon Hills, IL, USA) were programmed to generate ml/rev pulsatile perfusion, with 1-second-long fluctuations, and the perfusion pressures were recorded with a digital differential pressure manometer (Dwyer Instruments, Michigan City, IN, USA). The associated mean perfusion pressures were 97.21 mmHg, 87.56 mmHg, and 25.46 mmHg for the respective flow rates of 650 ml/min, 500 ml/min, and 200 ml/min. With these settings, scaffolds were subjected to constant physiologically normal (500 and 650 ml/min) and abnormal (200 ml/min) blood flow rates, and variations in renal blood flow patterns at three time points were recorded across 24 hours.

**Figure 5.**
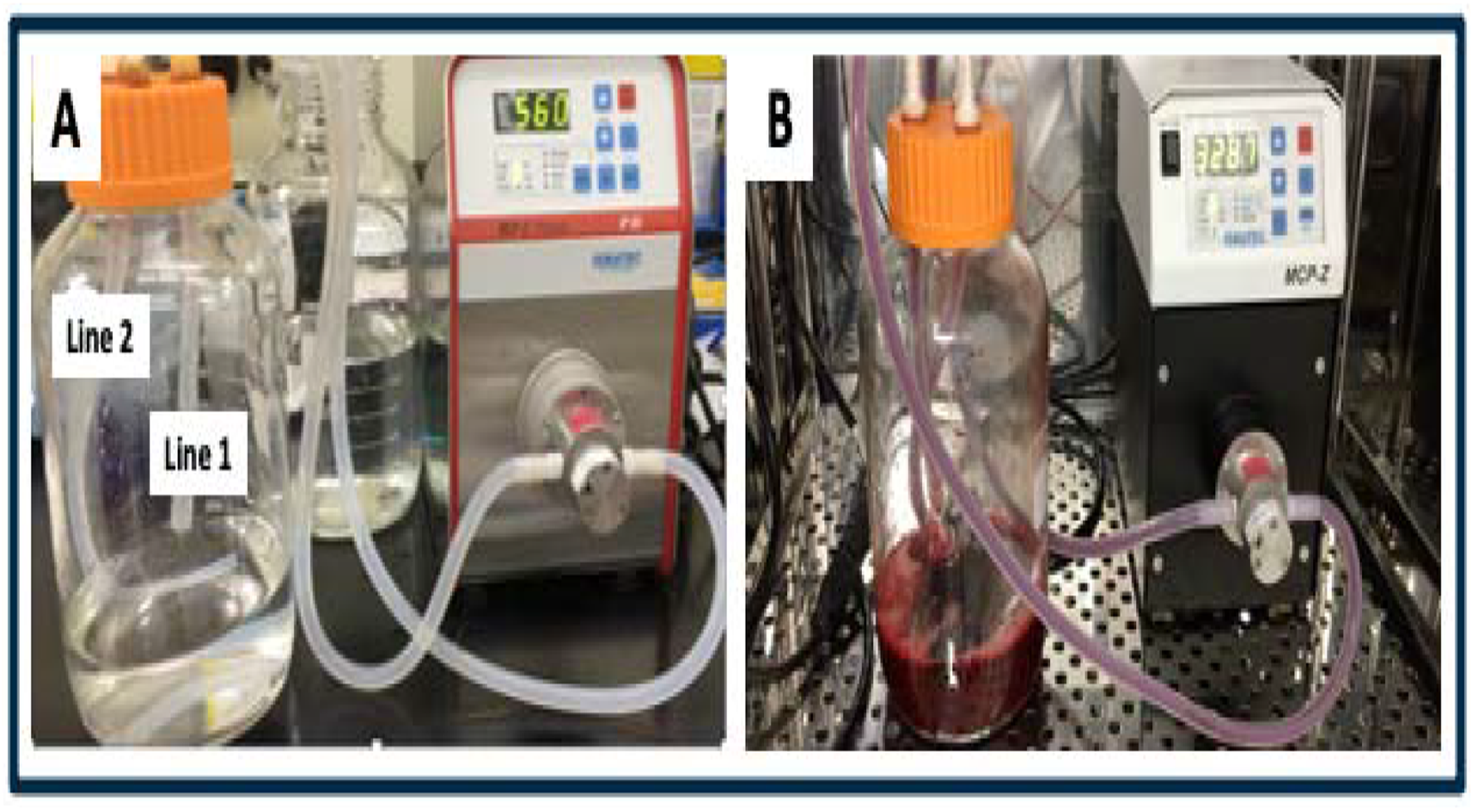
Images of the bioreactor model used to perfuse decellularized scaffolds with whole blood. (A) An image of the bioreactor system that was filled with sterilized PBS to flush the system initially. After this, the PBS was replaced by pig whole blood (B), and decellularized kidneys were suspended in the bioreactor by attaching the cannulated renal artery to line 1, while line 2 was left open to act as a venous reservoir to facilitate blood circulation.

### Statistical Analysis

Time-dependent drops in venous outflow were expressed as means ± standard deviation. The Student t-test with p < 0.05 level of significance was used to investigate the variations in venous outflow among the three groups.

## Data Availability

The datasets generated during and/or analyzed during the current study are available from the corresponding author on reasonable requests.

## Acknowledgments

The author would like to acknowledge Dr. Joao Paulo Zambon, Dr. James Yoo, and Dr. In Kap Ko at the Wake Forest Institute for Regenerative Medicine for supporting the initial formulation of this project. The author also wishes to thank Dr. Ali Khraibi, Dr. Dietrich Lorke, Dr. Ovidiu Baltatu, Ms. Khadija Al Hammadi, and Mr. Dawood Syed at Khalifa University of Science and Technology, and Dr. Gabriel Finkelstein at the University of Colorado for reviewing the manuscript.

